# Coupling CRISPR Scanning with Targeted Chromatin Accessibility Profiling using a Double-Stranded DNA Deaminase

**DOI:** 10.1101/2024.12.17.628791

**Authors:** Heejin Roh, Simon P. Shen, Yan Hu, Hui Si Kwok, Allison P. Siegenfeld, Ceejay Lee, Marc Anthony Zepeda, Chun-Jie Guo, Shelby A. Roseman, Vijay G. Sankaran, Jason D. Buenrostro, Brian B. Liau

## Abstract

Genome editing enables sequence-function profiling of endogenous cis-regulatory elements, driving understanding of their mechanisms and the development of gene therapies. However, these approaches cannot be combined with direct scalable readouts of chromatin structure and accessibility across long single-molecule chromatin fibers. Here we leverage a double-stranded DNA cytosine deaminase to profile chromatin accessibility at high depth and resolution at endogenous loci of interest through targeted PCR and long-read sequencing, a method we term targeted deaminase-accessible chromatin sequencing (TDAC-seq). Powered by high sequence coverage at targeted loci of interest, TDAC-seq can be uniquely integrated with CRISPR perturbations to enable the functional dissection of cis-regulatory elements, where genetic perturbations and their effects on chromatin accessibility are superimposed on the same single chromatin fiber and resolved at single-nucleotide resolution. We employed TDAC-seq to parse CRISPR edits that activate fetal hemoglobin in human CD34+ hematopoietic stem and progenitor cells during erythroid differentiation as well as in pooled CRISPR and base editing screens tiling an enhancer controlling the globin locus. Together, TDAC-seq enables high-resolution sequence-function mapping of single-molecule chromatin fibers by genome editing.

## Introduction

Cis-regulatory elements (CREs) are DNA sequences that function as switches to control the expression of genes.^1^ CREs often densely cluster in linear sequence space and contain finer-scale elements that together control transcription factor (TF) binding, chromatin accessibility, and downstream expression of associated genes.^2^ Methods that profile chromatin accessibility have provided high-resolution views of the function and actuation of CREs.^3–14^ Alternatively, CRISPR genome-editing technologies have enabled the systematic perturbation and fine mapping of endogenous CREs. For instance, high-resolution CRISPR mutational scanning of the *BCL11A* enhancer^15^ and *HBG1/2* promoter regions^16^ followed by readout of fetal hemoglobin (HbF) levels has provided key insights into specific elements and TF motifs that control HbF expression, ultimately culminating in breakthrough therapies to treat Sickle Cell Disease.^17–19^ Understanding how genome editing affects chromatin accessibility is essential for elucidating the molecular mechanisms that link genetic alterations to downstream phenotypic outcomes.

Technologies to jointly map and perturb CREs are needed and will drive basic understanding of DNA regulatory logic and development of gene therapies.^8–14^ Whereas traditional short-read measurements rely on fragmenting bulk populations of chromatin fibers,^3–7^ long-read sequencing formats retain information about crosstalk and interactions across a single chromatin fiber and enables synchronous readout of genetic variants and their effect on chromatin accessibility.^20^ However, these long read-compatible methods require higher DNA input amounts because they typically rely on methyltransferases to ‘stencil’ nucleosome and TF positioning, which are detected by sequencing the unnatural, methylated bases directly and are thus challenging to combine with PCR-based amplification. These limitations restrict their ability to study specific high-value loci at high coverage (>1000×), preventing integration with pooled CRISPR-Cas9 or base editor mutational scanning to measure the functional effects of genetic variants at scale. To address these limitations, we leverage DddA, a double-stranded DNA cytidine deaminase,^21^ to profile chromatin accessibility of targeted genomic loci using long-read sequencing after PCR amplification (**Figure 1A**). We term this method **<u>T</u>**argeted **<u>D</u>**eaminase-**<u>A</u>**ccessible **<u>C</u>**hromatin **<u>seq</u>**uencing (**TDAC-seq**). We leverage TDAC-seq with CRISPR perturbations to comprehensively read out editing outcomes with single-nucleotide resolution, enabling us to simultaneously map their direct impact on chromatin accessibility for hundreds of unique genotypes in a massively parallel pooled format.

**Fig. 1.**
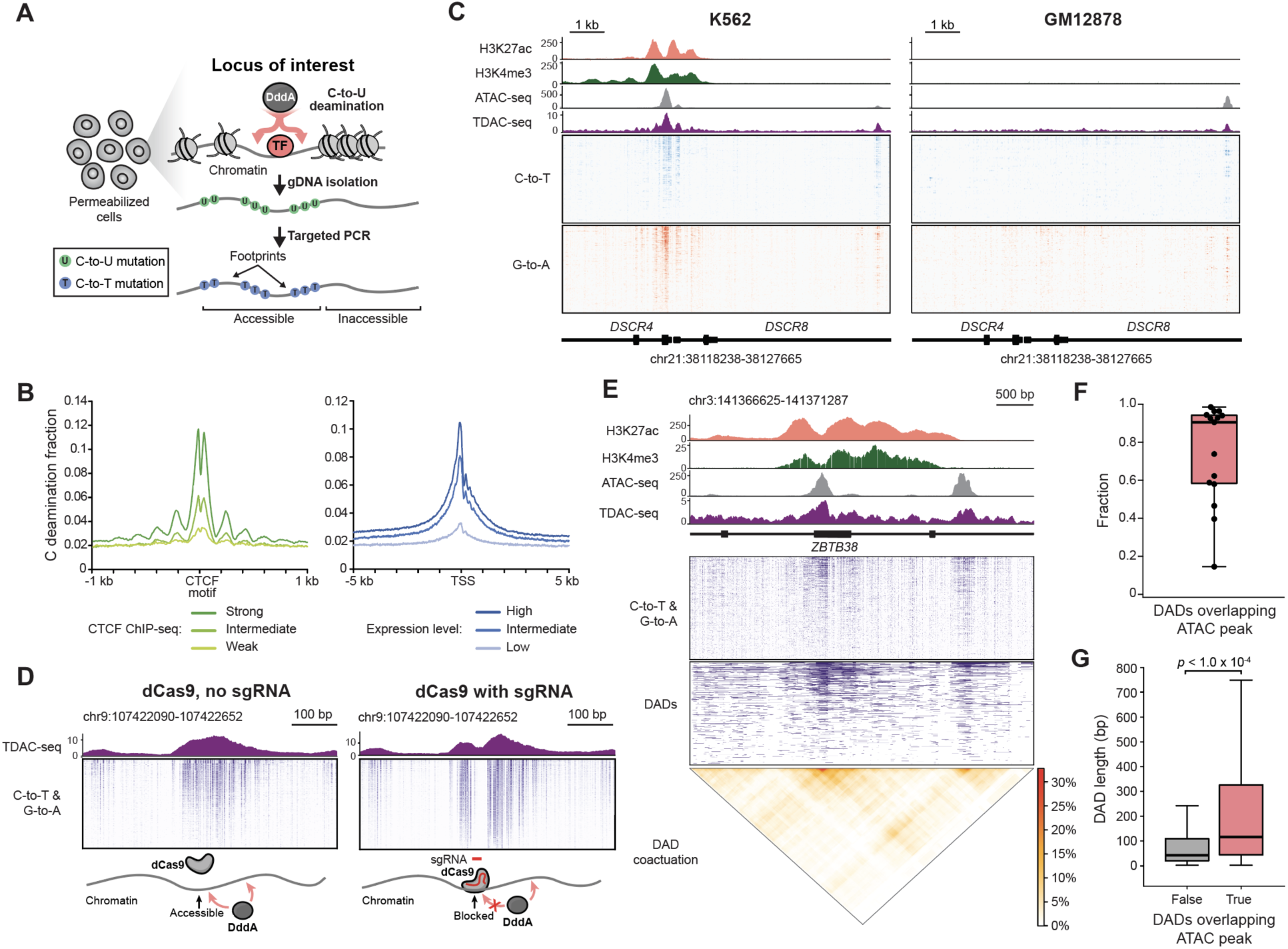
TDAC-seq detects chromatin accessibility across kilobase regions by targeted PCR. **A.** Schematic of TDAC-seq to map chromatin accessibility along targeted genomic loci using a double-stranded DNA cytidine deaminase. **B.** Aggregate profile plot showing distribution of cytidine deamination fractions (y axis) around CTCF binding sites (left) and transcription start sites (TSS, right) from whole-genome sequencing (WGS) of DddA11-treated K562 cells. **C.** TDAC-seq and chromatin tracks at the DSCR4/8 locus across a 9.4 kb region in K562 and GM12878 cells, showing cell-type-specific chromatin accessibility. TDAC-seq signal represents the average number of DddA11 mutations in a 50-bp window. C-to-T edits and G-to-A edits are depicted in red and blue, respectively. **D.** TDAC-seq tracks showing the footprint of dCas9 protein after introducing dCas9•sgRNA in MOLM-13 cells. C•G-to-T•A edits are depicted in purple. **E.** Top: genome tracks showing chromatin profiles, TDAC-seq, and DADs identified for each single DNA molecule at the ZBTB38 locus. Bottom: heatmap showing DAD co-activation, where color indicates the percentage of reads where both positions are open on the same DNA molecule. **F.** Box plot showing the fraction of DADs overlapping ATAC peaks in each of 15 TDAC-seq datasets. Box plots show the median and interquartile range, with whiskers extending to the minimum and maximum values. **G.** Box plots showing the distribution of DAD lengths across all reads in 15 TDAC-seq datasets, grouped according to overlap with ATAC peaks. Box plots show the median and interquartile range, with whiskers extending to the minimum and maximum values, excluding outliers that are beyond 1.5× the interquartile range. Welch’s t-test was used to calculate p-value.

### TDAC-seq for mutational profiling of chromatin accessibility on long, targeted loci

We sought to develop a method for employing DddA to mutationally profile chromatin accessible regions on long DNA molecules via C to U conversion, which preserves DNA integrity and creates C•G-to-T•A mutations after PCR amplification (**Figure 1A**). For this purpose, we employed DddA11, an evolved variant of the deaminase DddA that possesses relaxed sequence bias.^22^ Briefly, cell nuclei were isolated, permeabilized, and treated with purified DddA11. We first performed whole-genome sequencing of DddA11-treated K562 cells using Oxford Nanopore Technologies to assess C•G-to-T•A mutation rates and their positions. Consistent with prior reports,^22^ DddA11 preferentially deaminates cytidine preceded by a 5’-thymidine or 5’-cytidine (**Figure S1A**). Across the whole genome, C•G-to-T•A mutations were enriched at DNase I hypersensitive sites, CTCF sites, and active promoters, indicating that DddA11 preferentially deaminates accessible chromatin with a periodic mutation distribution consistent with strong nucleosome phasing at these regions (**Figure 1B**, **Figure S1B**). Altogether, these results demonstrate the ability of DddA11 to mark accessible chromatin by mutational profiling.

Since chromatin accessibility profiling with DddA11 does not cleave the DNA molecule but superimposes a mutational signature, our method is compatible with targeted PCR amplification and long-read sequencing technologies, allowing us to enrich specific genomic loci of interest with high coverage. To this end, we PCR-amplified fifteen genomic loci of interest after DddA11 treatment and sequenced the amplicons (∼3 to 11.5 kb in length) by nanopore sequencing. We then deduplicated reads using DddA11-induced mutations as unique molecular identifiers, assuming there is a very low likelihood of observing kilobase-sized molecules with identical distributions of stochastic C•G-to-T•A mutations (see Methods). This workflow results in TDAC-seq signal at the targeted loci of interest at high coverage, which were concordant with ATAC-seq data (**Figure 1C, Figure S1C**). Furthermore, dynamic changes to chromatin accessibility caused by dCas9 binding could be robustly detected by TDAC-seq (**Figure 1D**), highlighting the sensitivity of our approach and ability to detect protein occupancy.

Because TDAC-seq provides single-molecule readouts of chromatin accessibility across long DNA reads, we next investigated whether it could reveal correlations between accessible sites on single chromatin fibers that are otherwise obscured in bulk population average measurements (e.g., ATAC-seq). We examined the patterns of DddA11-induced mutations across individual DNA molecules and, as in previous approaches,^9,13^ observed long stretches of mutated bases corresponding to continuous open chromatin regions, which we refer to as deaminase-accessible DNA sequences (DADs) (**Figure 1E**). To correct for the variable density of DddA11-editable bases in different loci, we called DADs using a hidden Markov model with emission probabilities corresponding to the intrinsic sequence bias of the enzyme (**Figure S1D**). As validation of the DAD calling approach, across all sequencing reads in the fifteen loci of interest, a median of 90.4% of DADs overlapped with ATAC-seq peaks, and DADs within ATAC-seq peaks were significantly longer than those outside (**Figure 1F,G**). Next, we calculated per-read correlation of TDAC-seq signal across the respective loci. Consistent with prior reports,^9,23^ our analysis revealed nearby distal TDAC-seq accessible sites were correlated and open together across single chromatin fibers, highlighting the strength of long-read sequencing in detecting co-occurring accessibility (**Figure 1E, Figure S1E**). Altogether, these results establish TDAC-seq as a robust method to mutationally profile chromatin accessibility at target genomic loci of interest at high coverage.

### TDAC-seq reveals chromatin dynamics induced by CRISPR perturbations

In addition to measuring chromatin accessibility, we further tested the ability of DddA11 mutations to footprint nucleosomes and TFs, where the depletion of deamination-induced mutations within accessible chromatin could indicate protein binding. We calculated such depletion in varying footprint window sizes (1-99 bp) to detect footprints of objects with diverse sizes, where TFs were detected using smaller windows and nucleosomes are detected with larger windows (**Figure S2A,B**).^24^ To ensure that mutation rates reflect chromatin accessibility, the intrinsic sequence bias of DddA11 was measured and normalized using a bias model generated from deproteinized, naked gDNA treated with DddA11. Whole-genome sequencing of DddA11-treated nuclei revealed that different TFs display distinct patterns and intensities of footprints and associated nucleosome positioning. The aggregate TF footprint scores correlate with DNase-seq footprint scores,^25^ particularly for TFs known to exhibit strong footprinting in DNase-seq and ATAC-seq (**Figure S2C**). Nucleosome footprints from TDAC-seq align well with previously obtained MNase-seq data (**Figure S2D,E**).^26^ In addition, TDAC-seq on a region containing a CTCF binding site revealed the presence of strong CTCF footprints across single molecules (**Figure 2A, Figure S2D**). Altogether, TDAC-seq leverages DddA11 mutations to detect both nucleosome positioning and binding of TFs that produce strong footprints, providing single-read resolution within specific genomic loci that are PCR amplified.

**Fig. 2.**
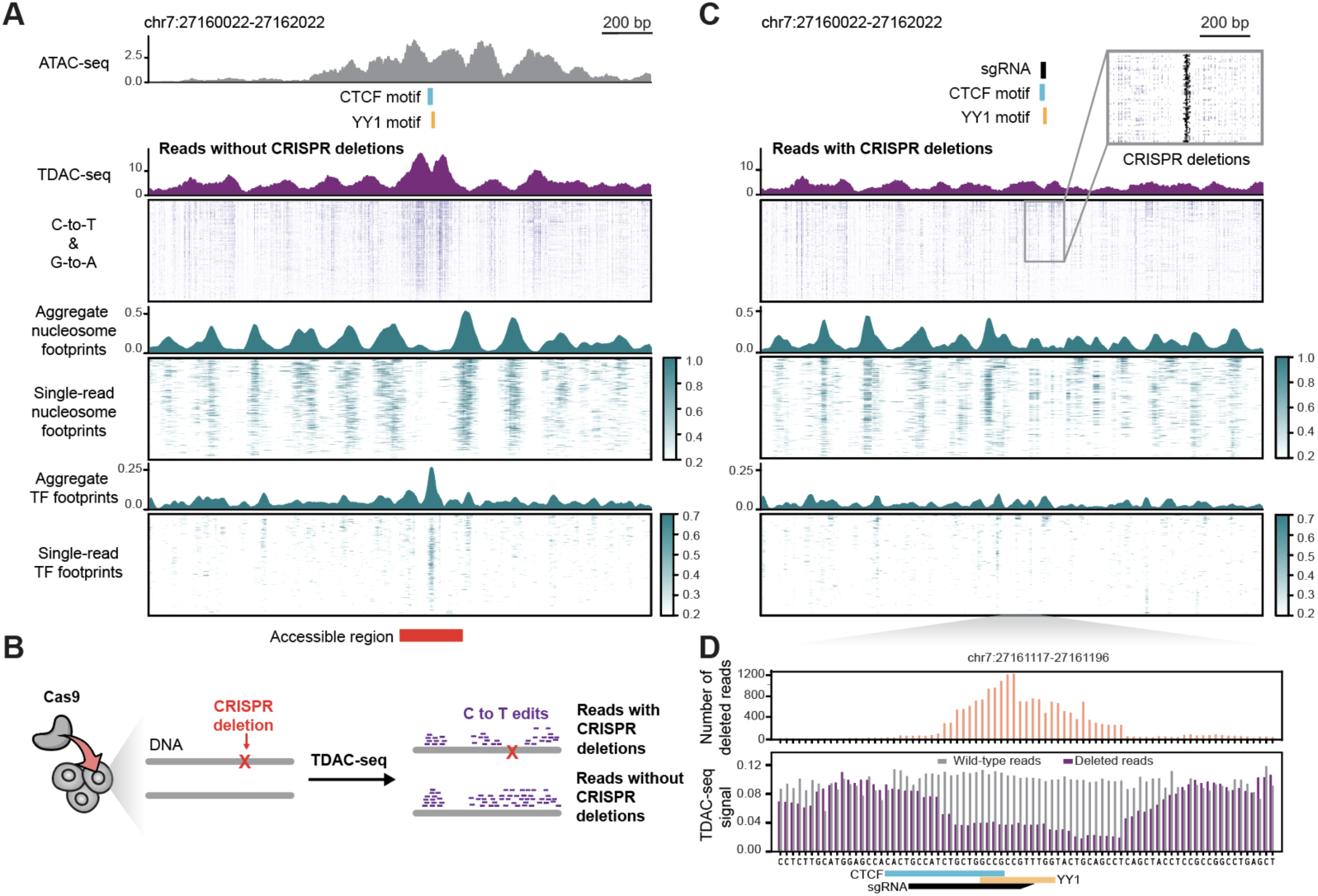
TDAC-seq measures CRISPR perturbation and chromatin structure from the same single DNA molecule. **A.** Genome tracks showing ATAC-seq, TDAC-seq, and footprint scores at the indicated CTCF binding site in MOLM-13 cells. The footprint window radii are 50 and 25 bp for nucleosome and TF footprints, respectively. **B.** Schematic of how TDAC-seq detects both CRISPR deletions and chromatin accessibility simultaneously from the same single DNA molecule, allowing partitioning of reads with and without CRISPR deletions. **C.** Genome tracks showing TDAC-seq and footprint scores from DNA reads containing CRISPR deletions near the sgRNA binding site, which overlaps the CTCF and YY1 motifs shown in Figure 2A. Magnified inset (top) highlights the CRISPR deletions (black). **D.** Top: Bar plot showing the number of reads (*y* axis) containing deletions within a 25 bp window centered on each nucleotide position (*x* axis) andno deletions observed outside this window. Bottom: Purple bars represent the average DddA mutation rate (*y* axis) from reads with deletions, while grey bars represent the mutation rate from reads without deletions, randomly sampled to match the number of reads with deletions. DddA mutation rates are calculated over the accessible region (red bar in Figure 2A) excluding each 25-bp deletion window.

Next, we tested whether TDAC-seq could distinguish changes in chromatin accessibility and footprinting following CRISPR perturbation. CRISPR-Cas9-mediated disruption of a CTCF binding motif in the *HOXA* locus was previously demonstrated to disrupt CTCF binding and decrease chromatin accessibility within the region.^27^ A sgRNA targeting this position and Cas9 were delivered into MOLM-13 cells, which was followed by TDAC-seq at this region. We then deconvoluted CRISPR-edited versus non-edited reads as well as the superimposed DddA11 mutational signature simultaneously by nanopore sequencing (**Figure 2B**). Compared to non-edited reads, DNA strands where the CTCF motif was disrupted showed reduced chromatin accessibility as well as weaker nucleosome and TF footprints (**Figure 2A,C, Figure S2F**). Moreover, since TDAC-seq affords single-nucleotide resolution, we could scrutinize the CRISPR deletions to determine the minimal deletion necessary to reduce chromatin accessibility (**Figure 2D**). This analysis revealed that deletions spanning the CTCF and YY1 motifs displayed the largest differential TDAC-seq signal, consistent with the previously reported roles of CTCF and YY1 in regulating chromatin accessibility.^27–29^ Together, these results demonstrate how TDAC-seq can simultaneously detect both CRISPR perturbations and chromatin accessibility from a single DNA molecule at single-nucleotide resolution.

### High-resolution profiling of chromatin accessibility in the β-globin locus during erythroid differentiation of CD34+ HSPCs

Profiling chromatin accessibility of targeted loci of interest at high coverage from primary cells remains challenging due to limiting amounts of input gDNA, rendering it difficult to achieve sufficient sequencing coverage. Since DddA-induced mutations are uniquely compatible with targeted PCR amplification, we posited that TDAC-seq could enable the high-resolution readout of chromatin accessibility for CREs of interest in primary cells. Consequently, we tested if TDAC-seq could resolve changes in chromatin accessibility in the β-globin locus upon erythroid differentiation of CD34+ hematopoietic stem cell progenitor cells (HSPCs) (**Figure 3A**). During erythroid differentiation, *HBB* and *HBD* gain and lose chromatin accessibility, respectively,^30,31^ and TDAC-seq of an 11 kb amplicon spanning the promoters of these two genes could robustly detect these expected changes (**Figure 3B, Figure S3A**).

**Figure 3.**
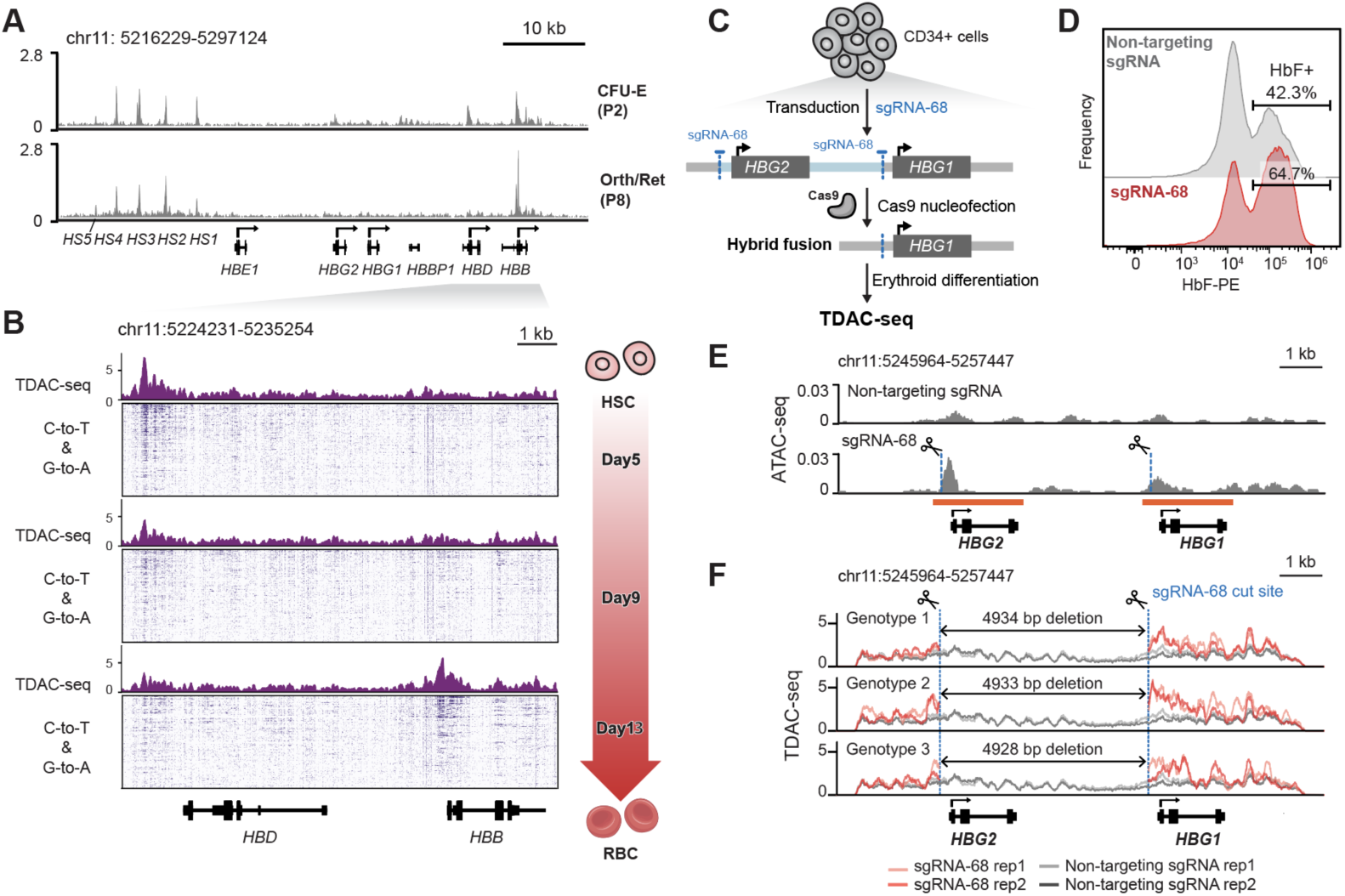
TDAC-seq detects chromatin accessibility dynamics at the β-globin locus in erythroid-differentiated HSPCs. **A.** ATAC-seq tracks showing the β-globin locus at indicated stages of CD34+ HSPC erythroid differentiation ^30^. **B.** Genome tracks showing chromatin accessibility changes measured by TDAC-seq at indicated stages of CD34+ HSPC erythroid differentiation at the *HBD-HBB* locus. **C.** Schematic of TDAC-seq combined with CRISPR-Cas9 targeting *HBG1/2* promoters in CD34+ HSPC cells. Cas9•sgRNA-68 targets two sites in the *HBG1/2* locus, which after cutting predominantly results in a hybrid fusion product containing an ∼5 kb deletion. TDAC-seq was conducted five days after inducing erythroid differentiation. **D.** Flow cytometry histograms of HbF levels (PE, *x* axis) in erythroid-differentiated HSPCs (day 5) treated with Cas9 and a non-targeting sgRNA (top, gray) or sgRNA-68 (bottom, red). **E.** ATAC-seq from erythroid-differentiated HSPC cells (day 5) treated with Cas9 and a non-targeting sgRNA or sgRNA-68. Although sgRNA-68 produces a hybrid fusion, ATAC-seq reads were aligned to both the *HBG1* and *HBG2* promoters due to homologous DNA sequences (orange lines). **F.** TDAC-seq of the three most abundant genotypes with different deletion sizes at the *HBG* locus from erythroid-differentiated HSPC cells (day 5) transduced with sgRNA-68, compared with non-targeting sgRNA control. TDAC-seq signals were aggregated from the top 10% of highly DddA-mutated reads. The blue dotted line indicates the sgRNA-68 cut sites.

Notably, the β-globin locus also encompasses the paralogous genes *HBG1* and *HBG2,* which control the expression of *γ*-globin during development, and have attracted interest for their therapeutic relevance but have been challenging to disentangle. Although typically silenced in adult blood, reactivation of *γ*-globin by mutations associated with hereditary persistence of HbF reduces the severity of Sickle Cell Disease.^15,18,32,33^ Due to this connection, the promoter regions of *HBG1* and *HBG2* have been targeted by CRISPR-based therapeutics in patient-derived CD34+ HSPCs to induce reactivation of *γ*-globin and HbF expression for the treatment of Sickle Cell Disease,^19,34^ This includes OTQ923, an autologous in ex vivo CRISPR-Cas9-edited CD34+ product, which ultimately progressed into patients.^19^ Intriguingly, sgRNA-68, the sgRNA component of OTQ923, targets the promoters of both *HBG1* and *HBG2* due to their high levels of sequence homology. Subsequent DNA repair and ligation of double-strand DNA break ends lead to a ∼5 kb deletion that excises *HBG2* entirely and creates a single hybrid gene with part of the *HBG2* promoter sequence fused to *HBG1* (**Figure 3C**). However, the impact of CRISPR edits on the locus structure and chromatin accessibility has been difficult to precisely measure by short-read-based approaches (i.e., ATAC-seq) because *HBG1* and *HBG2* are contiguous, duplicated genes spanning ∼7 kb, preventing the alignment of short reads to either specific gene region due to their high levels of sequence homology.

To determine whether our long-read approach could address these limitations, we conducted TDAC-seq on the *HBG1-HBG2* locus in CD34+ HSPC-derived erythroblasts, which were transduced with either a non-targeting sgRNA-control or sgRNA-68, electroporated with Cas9 protein, and induced to differentiate for 5 days **(Figure S3B**) — sufficient time to allow complete Cas9 turnover.^35^ We verified that Cas9 sgRNA-68 led to the creation of a large ∼5 kb deletion as the major product (**Figure S3C**), which upon differentiation resulted in robust upregulation of HbF (**Figure 3D)**. Due to the high sequence homology between the two promoters, short sequencing reads obtained from bulk ATAC-seq cannot distinguish them or detect these large deletions from the mixture of products resulting from Cas9 treatment (**Figure 3E**). By contrast, the large deletions could be clearly called for each TDAC-seq read, where the 4928-bp, 4933-bp, and 4934-bp deletions accounted for the vast majority of CRISPR mutational outcomes (**Figure S3D)**. Notably, chromatin accessibility could be measured simultaneously on reads containing these deletions, which displayed similarly elevated TDAC-seq signals in the promoter of the hybrid gene locus (**Figure 3F**), consistent with strong upregulation of HbF induced by sgRNA-68 and the notion that the deleted region may contain repressive elements.^34^ Altogether TDAC-seq enables a high-resolution view of the effects of OTQ923 per genotype on the *γ*-globin locus in primary CD34+ HSPCs upon differentiation.

### Integrating TDAC-seq with pooled CRISPR-mutational scanning

Given the high coverage afforded by TDAC-seq at targeted genomic loci of interest and the ability of our protocol to simultaneously detect CRISPR deletions and chromatin accessibility, we considered whether TDAC-seq could be combined with pooled CRISPR-mutational scanning to systematically dissect CREs to identify elements that control chromatin accessibility (**Figure 4A**). A pooled library consisting of 21 sgRNAs^36^ tiling hypersensitive site 2 (HS2) of the globin locus control region was transduced into K562 cells followed by electroporation of Cas9 and TDAC-seq after 6 days. Due to the stochasticity of DNA repair of the double-stranded breaks created by Cas9, we observed 541 unique CRISPR-induced deletion genotypes with sufficient sequencing coverage (>400 unique reads) to simultaneously measure their impact on accessibility (**Figure S4A,B**). We quantified C•G-to-T•A aggregate mutation rates per each genotype across the amplicon (**Figure 4B, Figure S4C**). DNA molecules bearing CRISPR deletions had variable TDAC-seq signal depending on the position and size of the CRISPR deletions. Notably, deletions attributed to sgRNA-10 and sgRNA-11 reside close to the center of the HS2 ATAC-seq peak and resulted in significantly reduced TDAC-seq signal. We clustered reads with these deletions to identify a shared minimal sequence whose deletion is necessary and sufficient for this effect (**Figure S4D**; see Methods). This analysis revealed a 10-bp sequence that corresponds to the AP-1 (JUN/FOS) motif, which has previously been demonstrated to be essential for activity of HS2.^37–39^ Reads where this motif is entirely deleted had significantly reduced DddA accessibility at HS2 compared to those that partially delete the motif or that do not perturb it (**Figure 4C**). Although CRISPR deletions will remove a small number of C•G base pairs that could have been edited by DddA, the number of DddA edits within these deletions is negligible relative to this effect size (**Figure S4E**). We also confirmed that individual introduction of sgRNA-11, as opposed to sgRNA-1 or a non-targeting control sgRNA, resulted in decreased ATAC-seq signal at HS2 as well as decreased expression of proximal globin genes (**Figure 4D,E**). While bulk ATAC-seq provides a bulk readout of the aggregate effects of a mixture of editing outcomes (**Figure S4F**), TDAC-seq can precisely deconvolute the effects of individual editing outcomes on chromatin accessibility.

**Figure 4.**
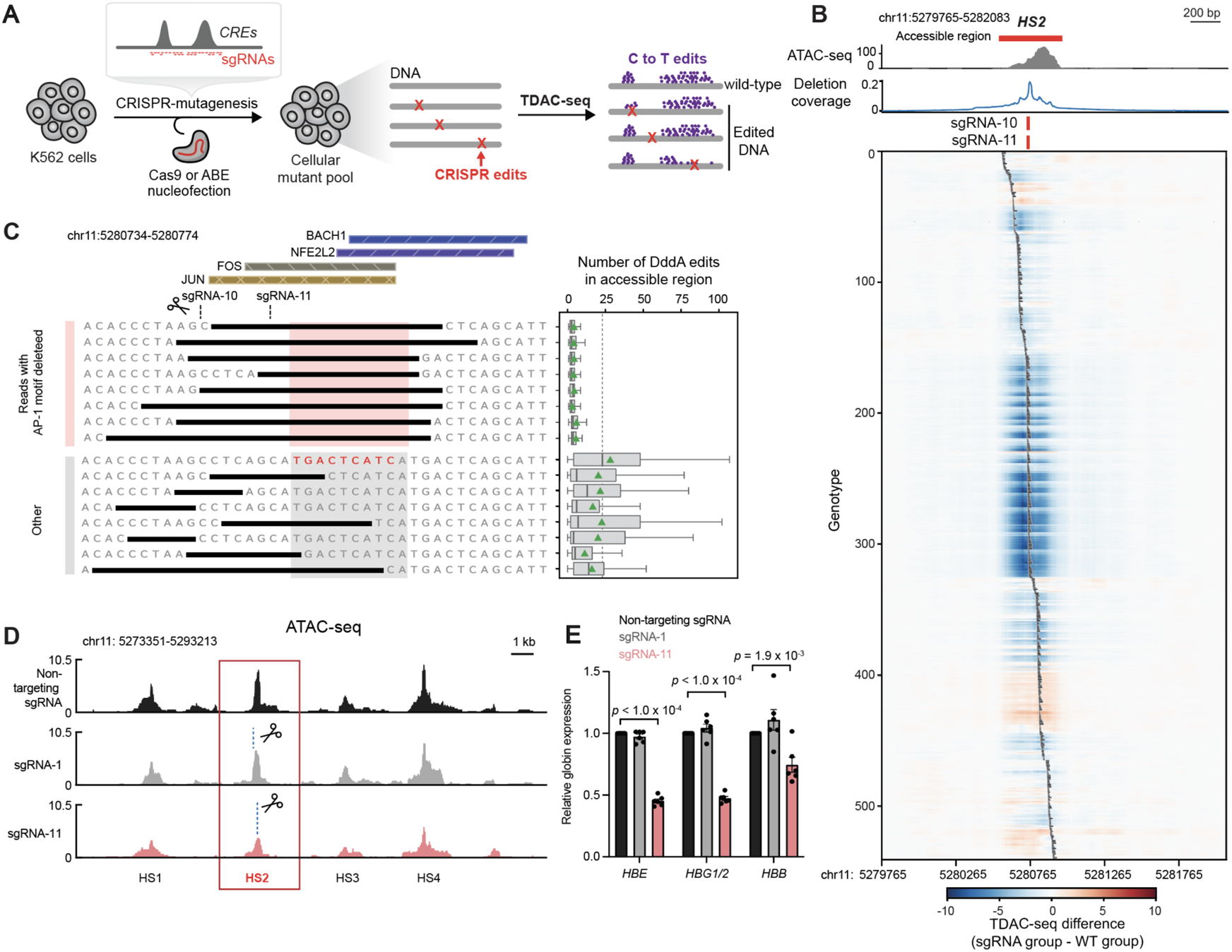
TDAC-seq can measure the impact of pooled CRISPR perturbations on chromatin accessibility in parallel. **A.** Schematic of TDAC-seq integrated with pooled CRISPR-mutational scanning of CREs to measure the effects of CRISPR perturbations on chromatin accessibility at target loci of interest. **B.** Top: wild-type K562 ATAC-seq and a line plot of the fraction of reads (*y* axis) containing a CRISPR deletion at the indicated position across the HS2 region (*x* axis). Bottom: Heatmaps of TDAC-seq using CRISPR-Cas9 cutting on the HS2 enhancer in K562 cells. Individual sequencing reads were grouped by 541 CRISPR deletion genotypes (y axis), with black bars indicating the CRISPR deletions on the HS2 region. For each genotype, TDAC-seq signal subtracted by wild-type signal is shown across the HS2 region (*x* axis). **C.** Genotypes of the most abundant edited alleles detected around the sgRNA-10 and sgRNA-11 cut sites (left) and the number of DddA edits (right) on those reads calculated in the accessible region (red bar in **Figure 4B**). Box plots show the median and interquartile range, with whiskers extending to the minimum and maximum values, excluding outliers that are beyond 1.5× the interquartile range. Mean is indicated by green triangles. Genotypes were clustered based on the 10-bp deletion window indicated with a red box. Edited alleles containing deletions that overlap the FOS/JUN (AP-1) motifs from Unibind ^47,50^ (key motif sequence labeled in red text) show the strongest decrease in chromatin accessibility. **D.** Genome tracks showing ATAC-seq signal (*y* axis) after individual transduction of a non-targeting sgRNA, sgRNA-1, and sgRNA-11 into K562 cells. The blue dotted lines indicate the sgRNA cut sites. **E.** Bar plot showing mRNA levels of *HBE, HBG1/2*, and *HBB* after individual transduction of sgRNA1 and sgRNA11 as measured by qRT-PCR. Relative globin expression fold-changes were calculated relative to K562 cells transduced with non-targeting sgRNA.

### TDAC-seq enables detection of cooperative ABE effects on CRE accessibility

Next, we sought to test the compatibility of TDAC-seq with finer-scale genetic perturbations afforded by base editors, specifically the adenosine base editor 8e (ABE8e) that yields A•T-to-G•C mutations.^40^ Purified ABE8e was electroporated into K562 cells transduced with the same HS2-tiling sgRNA library as before and TDAC-seq was then performed (**Figure 4A**). A•T-to-G•C mutations were strongly enriched at the predicted editing windows and target strands of the sgRNAs, consistent with editing by ABE (**Figure 5A**: upper panel, **Figure S5A,B**). Because many sgRNAs contain more than one adenosine within the ABE editing window, many reads stochastically had more than one A•T-to-G•C mutation. As a result, we observed 49 unique genotypes created by ABE with sufficient sequencing coverage (>100 unique reads) (**Figure S5C**). To evaluate the impact of ABE edits on TDAC-seq signal, we grouped reads by the position of ABE edits, computed the average TDAC-seq signal of each genotype, and compared with wild-type reads (**Figure 5A**, lower panel, **Figure S5D**). The impact of these single base edits on TDAC-seq signal was attenuated in comparison to deletions afforded by Cas9 nuclease, consistent with the generally weaker effect size of point mutations. Like with the deletions produced by Cas9 nuclease, ABE mutations targeting the center of HS2 exhibited the most significant reductions in TDAC-seq signal.

**Figure 5.**
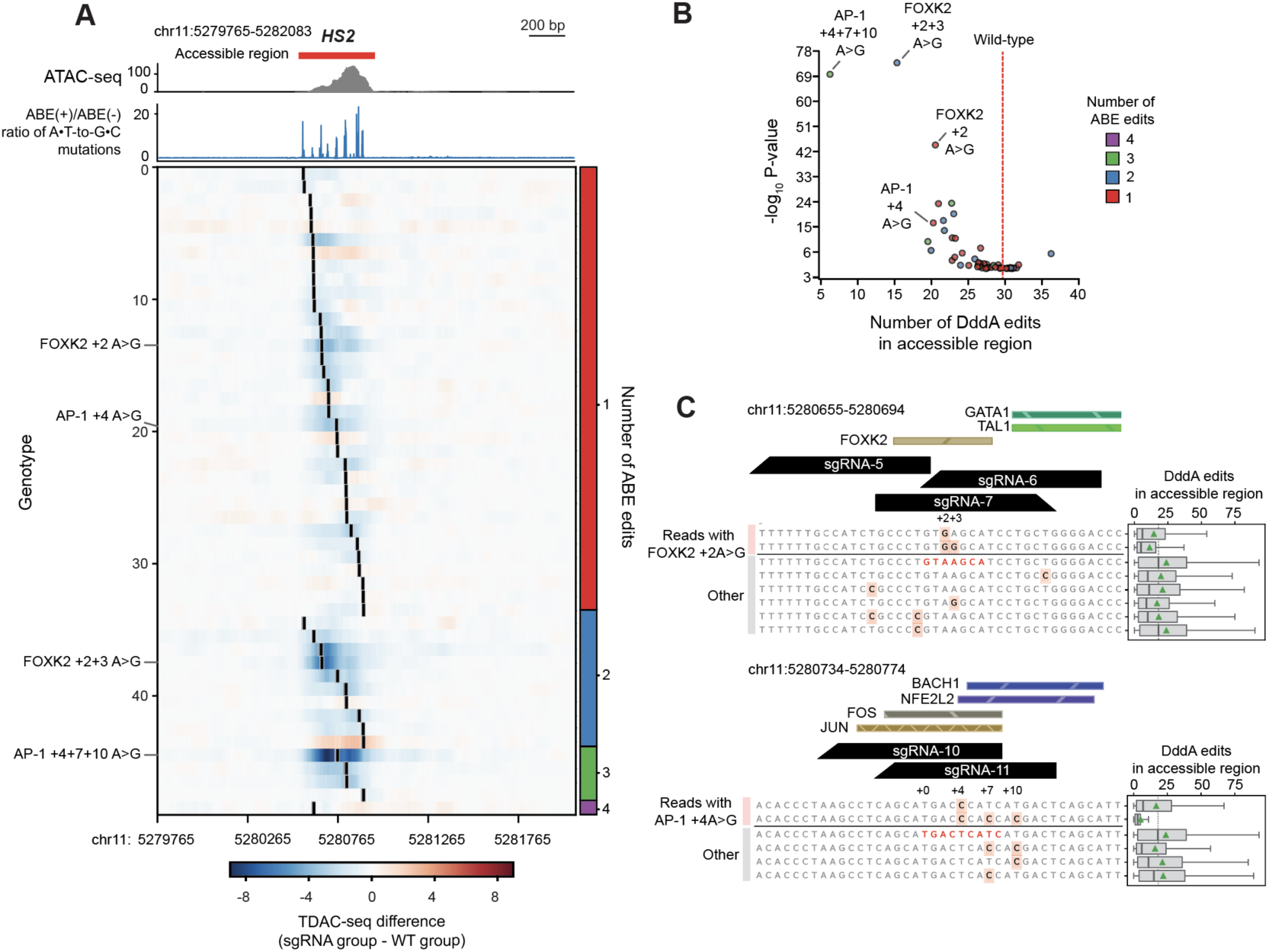
TDAC-seq can measure the impact of pooled ABE perturbations on chromatin accessibility across distinct genotypes. **A.** Top: Wild-type K562 ATAC-seq and a line plot showing the ratio of A•T-to-G•C edit coverage in ABE(+) versus ABE(-) samples across the HS2 region (*x* axis). Bottom: Heat maps of TDAC-seq using ABE on the HS2 enhancer in K562 cells. Individual sequencing reads were grouped by the positions of A•T-to-G•C base edits (*y* axis), with black lines indicating the edited locations on the HS2 locus (*x* axis). For each genotype, TDAC-seq signal subtracted by wild-type signal are shown across the HS2 region (*x* axis). Genotypes were grouped by the number of ABE edits (right). **B.** Volcano plot showing the number of DddA edits in accessible region (red bar in **Figure 5A**) and their -log10 *p*-value for each genotype. Wild-type reads have about 30 DddA edits in average (dotted red line). Each dot (genotype) is colored based on the number of ABE edits. Top hits are labeled by the positions of ABE edits within the TF motif. **C.** Box plots (right) showing the number of DddA edits for reads with the indicated genotype (left). A•T-to-G•C mutations are highlighted in orange. Genotypes are clustered by whether the labeled A•T base pair is mutated, and the most common genotypes with at least 1000 reads are shown. sgRNA binding sites and selected transcription factor binding sites from Unibind are shown above the sequences. The core motifs of FOXK2 (top) and FOS/JUN (AP-1) (bottom) are labeled in red text. DddA edits are counted over the accessible region (red bar in **Figure 5A**). Box plots show the median and interquartile range, with whiskers extending to the minimum and maximum values, excluding outliers that are beyond 1.5× the interquartile range. Mean is indicated by green triangles.

Since TDAC-seq can directly genotype base edits as opposed to indirectly inferring from sgRNA deconvolution, we considered whether the effects of multiple simultaneous base edits— which often occur when there are multiple adenosines within a sgRNA’s editing window—could be interrogated. TDAC-seq reads containing multiple edits (representing 18.6% of all sequencing reads) could be robustly detected, and as anticipated, tend to show stronger reduction in HS2 accessibility compared to reads with single base edits, highlighting cooperative effects (**Figure 5B, Figure S5E**). Notably, while the +2A>G mutation in the FOXK2 motif reduces HS2 accessibility and had the strongest effect among reads with a single base edit – consistent with the reported^41^ cooperativity between FOXK2 and AP-1 – combining it with +3A>G yielded even stronger effects (**Figure 5C**, upper panel). Likewise, the +4T>C mutation in the AP-1 motif was the second strongest individual base edit in our pool, and combining that mutation with the +7T>C and/or +10T>C mutations reduced accessibility even more than any of these mutations individually (**Figure 5C**, lower panel). Altogether, these findings show that TDAC-seq can be integrated with pooled CRISPR-mutational scanning at scale to dissect sequence-function relationships governing enhancer accessibility at single-nucleotide resolution.

## Discussion

In summary, TDAC-seq combined with CRISPR-mutational scanning provides targeted and scalable methods to fine-map the activity and sequence determinants of DNA cis-regulatory elements in high resolution across hundreds of editing outcomes. This integration is uniquely possible by leveraging a double-stranded DNA cytidine deaminase that mutationally profiles chromatin fibers in a manner compatible with PCR of the targeted loci, enabling the sequencing coverage and resolution at the locus of interest necessary for conducting a pooled CRISPR screen. Importantly, CRISPR edits can lead to diverse outcomes and potential combinatorial effects, and these details are not fully distinguished by only capturing sgRNA identity. Notably, TDAC-seq can precisely characterize heterogeneous editing outcomes directly while simultaneously profiling associated chromatin states, allowing us to dissect finer TF motifs controlling the accessibility of CREs at single-nucleotide resolution. More broadly, we anticipate that TDAC-seq will provide a high-throughput platform to systematically evaluate the impact of sgRNAs for CRISPR-Cas9 and base editing therapeutic strategies for the treatment of human genetic diseases.

## Acknowledgments

The authors would like to thank J. Nelson at the Bauer Core Facility of Harvard University for assistance with FACS and members of the Liau Lab and others for insightful comments and discussion on the manuscript, in particular F. Najm, E. Gaskell, and B. Bernstein. We thank R. Zhang, and M. Horlbeck for helpful discussions as well as technical and computational support. We thank B.Y. Mok and D. Liu for providing the DddA expression plasmids.

## Author contributions

HR, AS, and BL conceived the study. HR designed the experiments and performed protein purification, biochemical assays, and cell biology experiments with assistance from HSK. HR, SS, and YH performed computational analysis, with input from JB and BL. AS, SL, CJG, and SR provided experimental support with input from VS.HR, SS, YH, HSK, and BL wrote and edited the manuscript with input from all other authors. BL held overall responsibility for the study.

## Methods

### DddA11 expression and purification

DddA11 was cloned into pETDuet-1::*dddAt*ox-*dddI*A (a gift from D. R. Liu) by introducing six mutations in the DddA protein coding sequence (S1330I, A1341V, N1342S, E1370K, T1380I, T1413I).^21^ Mutant sequences were generated by PCR and introduced using NEBuilder HiFi DNA Assembly Master Mix (NEB E2621L). After obtaining the pETDuet-1::*dddA11t*ox-*dddI*A plasmid, a TEV cleavage site and an 8-amino-acid linker sequence, ‘GGGSGGGS,’ followed by the MBP tag, were inserted in-frame after the protein coding sequence.

The plasmid was transformed into BL21(DE3)pLysS *E.coli* cells. After plating, a single colony was cultivated in LB media with 100 mg/L ampicillin at 37 °C for 6 h. This culture was used to inoculate 12 L of LB broth at a 1:100 dilution, and the culture was grown at 37 °C until it reached an OD600 of 0.6. Expression was induced with 0.5 mM isopropyl β-D-thiogalactoside (IPTG), and the culture was incubated at 18 °C for 16 h. Cells were harvested by centrifugation at 4000g for 35 min, and each 3 L pellet was resuspended in 40 mL binding buffer (5 mM imidazole, 0.5 M NaCl, and 20 mM Tris–HCl, pH 7.9) containing 1 mg/mL lysozyme. The cells were sonicated at 60% amplitude for 10 min (10 s on, 10 s off) on ice, and the lysate was centrifuged at 12000 rpm for 40 min.

All protein purification steps were performed at 4 °C. A column was loaded with His60 nickel resin (Takara Bio), washed with 5 column volumes (cv) of Milli-Q water, and then with 6 cv of binding buffer. The supernatant was added to the charged resin and incubated on a rotator for 20 min. The supernatant was allowed to flow through the column, followed by sequential washes with 20 cv of binding buffer, 10 cv of wash buffer I (60 mM imidazole, 0.5 M NaCl, 20 mM Tris– HCl, pH 7.9), 5 cv of wash buffer II (100 mM imidazole, 0.5 M NaCl, 20 mM Tris–HCl, pH 7.9), and 7 cv of elution buffer (500 mM imidazole, 0.5 M NaCl, 20 mM Tris–HCl, pH 7.9, 1 mM DTT). Eluates were directly analyzed by Coomassie-stained SDS-PAGE. Fractions containing DddA11-MBP-DddI were collected and subjected to buffer exchange using a concentrator (Amicon Ultra-15 Centrifugal Filter Unit, 30 kDa MWCO) in refolding wash buffer (5 mM imidazole, 50 mM Tris– HCl pH 7.5, 500 mM NaCl, and 1 mM DTT). The concentrated protein was added to 350 mL of denaturing buffer (8 M urea, 5 mM imidazole, 50 mM Tris–HCl pH 7.5, 500 mM NaCl, and 1 mM DTT) and incubated for 36 h at 4 °C with shaking. A fresh Ni resin was washed with 50 cv of Milli-Q water and 10 cv of 8 M urea denaturing buffer. The urea buffer containing the eluted proteins was loaded onto a gravity-flow column, incubated for 20 min, and the flow-through was collected. Sequential washes were performed with 7 cv of denaturing buffer with decreasing urea concentrations (8 M, 6 M, 4 M, 2 M, 1 M), followed by a final wash with refolding wash buffer to remove residual urea. Proteins bound to the resin were eluted with 3 cv of wash buffer 1, followed by 1 cv of wash buffer 2, and finally 5 cv of elution buffer. Protein-containing fractions were identified by SDS-PAGE and subsequently subjected to two rounds of buffer exchange using a 30 kDa concentrator in loading buffer (20 mM Tris pH 8, 20 mM NaCl, 1 mM DTT). The concentrated samples were purified by anion exchange chromatography using fast protein liquid chromatography (FPLC) with a 5mL HiTrap Q HP column (Cytiva). The column was equilibrated in low-salt buffer (20 mM Tris–HCl pH 8, 1 mM DTT), and proteins were separated with a gradient from 100% low-salt buffer to 100% high-salt buffer (20 mM Tris–HCl pH 8, 1 M NaCl, 1 mM DTT) over 176 mL at a flow rate of 4 mL/min. Fractions were evaluated by SDS-PAGE. The DddA11 fractions, free of other contaminants, were collected, buffer-exchanged once with DddA storage buffer (20 mM Tris–HCl pH 7.5, 200 mM NaCl, 1 mM DTT, 5% (w/v) glycerol) using a 10 kDa concentrator, and stored at –80 °C.

### TDAC-seq DddA reaction

1× 10^6^ cells were harvested by centrifugation at 300×g for 5 min. The cell pellet was resuspended in 500 µL of PBS, gently pipetted to wash, and centrifuged again at 300×g for 5 min at 4 °C. The cell pellet was resuspended in 500 µL of cold lysis buffer, and incubated at room temperature for 10 min, followed by incubation at 4 °C for 5 min. After incubation, the cell pellet was centrifuged at 300×g for 5 min at 4 °C, and the supernatant was discarded. The cell pellet was then washed with 500 µL of wash buffer, centrifuged at 300×g for 5 min at 4 °C, and the wash buffer was removed. The activation buffer was prepared by adding purified DddA11-MBP. The cell pellet was resuspended in 50 µL of activation buffer by gentle pipetting, and the reaction mixture was incubated at 37 °C for 2 h with shaking at 300 rpm. Buffer volume was reduced proportionally if fewer cells were used, with reactions conducted for as few as 2× 10^5^ cells.

To isolate genomic DNA (gDNA), a 50 uL reaction was quenched by adding 5 µL of 10% SDS to a final concentration of 0.5% SDS, along with 35 µL of nuclease-free water and 10 µL of Proteinase K (10 mg/mL). The solution was incubated at 56 °C for 10 min. gDNA was isolated using a Zymo genomic DNA extraction kit (D4064), and the concentration was measured using a NanoDrop microvolume spectrophotometer.

### TDAC-seq library preparation

To minimize primer binding bias, we used primers containing Y and R in the TC and CC dinucleotide sequences, which are highly mutated sequence contexts by DddA11. 125-500 ng of gDNA was amplified per 25 µL PCR1 reaction (2× KOD One PCR Master Mix (TYB-KMM-101-5PK), 0.3 µM amplicon-specific forward primer, 0.3 µM amplicon-specific reverse primer). The PCR reaction was conducted with an amplicon size range up to 11.5 kb, under the following conditions: 30 cycles at 98°C for 10 sec, primer annealing temperature for 5 sec, and 68 °C for 15 min. For semi-nested PCR, the PCR2 reaction was set up by adding 1 µL of PCR1 product. PCR2 was conducted under the same conditions as PCR1 but with 3-8 cycles. 1 µL of PCR2 product was used in barcoding PCR3 with the nanopore PCR Barcoding Expansion 1-96 kit (EXP-PBC096), using 2× KOD One PCR Master Mix. This reaction was run for 12 cycles under the following conditions: 98 °C for 10 sec, 60 °C for 5 sec, and 68 °C for 15 min. To increase PCR yield, PCR1 or 2 products could be purified using 1.2× Mag-Bind® TotalPure NGS Kit (Omega Bio-Tek M1378-01), with the entire eluate used for the next PCR. The final amplicons were purified via DNA gel extraction using the Zymoclean Gel DNA Recovery Kit (D4001) and quantified using a Qubit fluorometer. Barcoded amplicons were pooled in desired ratios, and final nanopore libraries were prepared following the Ligation sequencing V14 (SQK-LSK114), followed by sequencing on a PromethION Flow Cell (R10.4.1) with super accuracy (SUP, v4.3.0) base calling model.

### DddA11-whole genome sequencing

gDNA extracted after the DddA reaction was fragmented to 10 kb size using Covaris G-tubes (Covaris 520079), and DNA concentration was measured with a Qubit fluorometer. Nanopore DNA libraries were then generated following the Ligation Sequencing V14 – PCR Barcoding (SQK-LSK114 with EXP-PBC096) protocol from Oxford Nanopore Technologies. Briefly, after end repair and barcode adapter ligation, barcoding PCR was performed with 2× KOD One PCR Master Mix, under the following conditions: 16 cycles of 98 °C for 10 sec, 60 °C for 5 sec, and 68 °C for 15 min. The PCR product was purified using 0.4× AMPure XP Beads, quantified using a Qubit fluorometer, and pooled with all barcoded libraries in desired ratios. A total of 1 µg of pooled barcoded libraries was prepared, and final nanopore libraries were completed following the Ligation Sequencing V14 protocol (SQK-LSK114) with end-prep and adapter ligation, followed by sequencing on a PromethION Flow Cell (R10.4.1) with super accuracy (SUP, v4.3.0) base calling model.

### Generating deproteinized DNA data

gDNA was extracted from K562 cells using the QIAamp DNA Blood Kit (Qiagen 51104), yielding deproteinized DNA. A 12.5 µL DddA reaction was prepared as described above, using 5 µg of purified gDNA. The reaction was incubated for 2 h, with shaking at 300 rpm at 37° C in a ThermoMixer. To quench the reactions, 1.25 µL of 10% SDS was added to each reaction to achieve a final concentration of 0.5% SDS, followed by 8.75 µL of nuclease-free water and 2.5 µL of Proteinase K (10 mg/mL). The mixture was incubated at 56 °C for 10 min, and gDNA was subsequently isolated using the Zymo Genomic DNA Extraction Kit. The edited DNA was amplified for eight distinct amplicons, and the DNA library was prepared and sequenced as previously described.

### Cell culture and lentiviral transduction

K562 was obtained from ATCC (CCL-243); HEK 293T was a gift from Bradley E. Bernstein; GM12878 was a gift from Xiaowei Zhuang; and MOLM-13 was a gift from Matthew D. Shair. All cell lines were authenticated by Short Tandem Repeat profiling (Genetica) and routinely tested for mycoplasma (Sigma-Aldrich). K562 and MOLM-13 were cultured in RPMI-1640 (Gibco) supplemented with 10% FBS (Peak Serum) and GM12878 were cultured in RPMI-1640 (Gibco) supplemented with 15% FBS. HEK293T and were cultured in DMEM (Gibco) supplemented with 10% FBS. All media were supplemented with 100 U/mL penicillin and 100 µg/mL streptomycin (Life Technologies). All cell lines were cultured in a humidified 5% CO2 incubator at 37 °C. Lentivirus was produced by co-transfecting HEK293T cells with pCMV-VSV-G (a gift from Bob Weinberg; Addgene plasmid #8454), psPAX2 (a gift from Didier Trono; Addgene plasmid #12260) and the transfer vector plasmid encoding gene of interest. Transfections were performed using Lipofectamine 3000, following the manufacturers’ protocols. Media was exchanged after 6-12 h and the viral supernatant was collected 48 h post-transfection and filtered (0.45 µm). Transduction was carried out by mixing the virus with cells with 8 µg/ml polybrene for K562 and 5 µg/ml polybrene for MOLM-13 then spinfected at 1,800 x *g* for 90 min at 37 °C. After 48 h post-transduction, media was changed and puromycin (Thermo Fisher Scientific) selection was carried out for 4 days.

### Generation K562 cell lines expressing dCas9 with sgRNA

The pHR-SFFV-dCas9-BFP plasmid was subcloned from pHR-SFFV-dCas9-BFP-KRAB (gift from Stanley Qi and Jonathan Weissman, Addgene #46911). K562 cells were transduced with lentiviral particles carrying the pHR-SFFV-dCas9-BFP plasmid as described above and cells with BFP positive cells were sorted on a MoFlo Astrios EQ cell sorter. The sgRNA sequence targeting the *HEK3* locus, GGCCCAGACTGAGCACGTGA, was cloned into pLentiGuide-Puro (gift from Feng Zhang, Addgene plasmid #52953). Lentiviral particles carrying the plasmids were generated as described above, and K562 dCas9-BFP cells were transduced and subsequently selected with puromycin (2 µg/mL).

### Human CD34+ HSPCs culture and three-phase erythroid differentiation

Human CD34+ HSPCs from mobilized peripheral blood of healthy adults were obtained from the Cooperative Center of Excellence in Hematology at the Fred Hutchinson Cancer Research Center. The primary HSPCs were thawed into a maintenance medium consisting of a StemSpan II base (StemCell Technologies), CC100 (StemCell Technologies), 50 ng/mL human TPO (Pepro Tech), 35nM UM171 (StemCell Technologies) and 1% penicillin/streptomycin (Life Technologies) and 1% of L-Glutamine (Life Technologies).

After the maintenance phase, primary human HSPCs were differentiated using the three-phase culture system previously described.^42,43^ First, a base erythroid medium was created by supplementing IMDM (Gibco) with 2% human AB plasma (SeraCare), 3% human AB serum(Life Technologies), 3 U/mL heparin, 10 μg/mL insulin, 200 μg/mL holo-transferrin, and 1% penicillin/streptomycin. From days 1-7 in erythroid media, this base medium was further supplemented with 3 U/mL EPO, 10 ng/mL human SCF, and 1 ng/mL IL-3. From days 7-12, this base medium was further supplemented with 3 U/mL EPO and 10 ng/mL human SCF. After day 12, the base medium was supplemented with 1 mg/mL total of holo-transferrin and 3 U/mL of EPO.

### HBG1/2 Cas9 deletions in CD34+ HSPCs with erythroid differentiation

The sgRNA-68 sequence targeting HBG promoter is ACTGAATCGGAACAAGGCAA. Oligos containing the sgRNA sequences were cloned into pLentiGuide-Puro (Addgene plasmid #52953). Lentiviral particles carrying the plasmids were generated as described above. Cell supernatant was collected and filtered through a 0.45 mM membrane and concentrated using Lenti-X Concentrator (Takara Bio, 631232).

Human CD34+ cells were cultured in hematopoietic stem cell (HSC) expansion media consisting of StemSpan SFEM II (StemCell Technologies, 02690) supplemented CC100 cytokine cocktail (StemCell Technologies, 02690), 50 ng/mL human TPO (PeproTech, 300-18), 1% L-Glutamine (Gibco), 1% penicillin/streptomycin (Gibco), and 35 nM UM171. After a one-day maintenance phase, lentivirus was pre-coated on plates with Retronectin (Takara Bio, T110A) and cells were transduced by spinfection at 2,000 rpm for 1.5 h at 37°C and cultured for 3 days.

To generate double stranded DNA breaks with sgRNA-68, Alt-R Cas9 protein (IDT) was transfected into cells via electroporation using NEON system (Thermo Fisher Scientific) with the following conditions: 1600 V, 10 ms, 3 pulses. Following electroporation, cells were transferred into pre-warmed HSC media and placed in culture for 2 days.

For erythroid differentiation, CD34+ cells were differentiated in vitro toward the erythroid lineage using an adaptation of the culture method.^15^ Briefly, cells were cultured in erythroid differentiation medium (EDM), consisting of Iscove’s Modified Dulbecco’s Medium (IMDM) (Gibco), 330 mg/ml human holotransferrin (Sigma, T4132), 10 mg/ml recombinant human insulin (Sigma, I9278), 2 IU/ml heparin (Sigma, H3149), 5% human AB serum (Research Products Internation, H68250), 2.5 U/ml human erythropoietin (PeproTech, 100-064), 1% glutamine (Gibco) and 1% penicillin/streptomycin (Gibco). EDM was further supplemented with 1.38 mM hydrocortisone (StemCell Technologies, 07925), 100 ng/ml human SCF (PeproTech, 300-07), 5 ng/ml human IL-3 (PeproTech, 200-03) and 1 mg/ml puromycin (Gibco) for cell selection. Cells were grown in humidified incubators at 37° C and 5% CO2. Cells were harvested for cell staining, ATAC-seq, and TDAC-seq on Days 5 post-differentiation induction.

### Staining for flow cytometry

Human CD34+ cells were stained at day 5 of erythroid differentiation for 2-phase culture system. For staining of CD71 and CD235a surface markers, cells were washed once with PBS and stained with 1:50 dilution of anti-CD71-BV711 (BD Biosciences, BDB5637670) and anti-CD235a-PE (eBioscience, 12-9987-82) for 30 min at 4 °C. For intracellular staining of HbF, cells were washed, fixed with fixation buffer (Biolegend, 420801) and permeabilized with intracellular staining permeabilization wash buffer (Biolegend, 421002). Cells were then incubated with 1:40 dilution of anti-HbF-PE for 20 min at 4 °C. After staining, the cells were washed twice with wash buffer, resuspended in staining buffer. Data acquisition was performed on ACEA Novocyte flow cytometer (Agilent) using NovoExpress software (version 1.5.6). Data were analyzed with FlowJo (version 10.10).

### ABE8e purification

ABE8e was purified as previously described.^40^ Briefly, the plasmid (Addgene plasmid #161788) was transformed into BL21(DE3)pLysS *E.coli* cells, and cultures were maintained in TB media with 25 mg/mL kanamycin. Following saturated overnight culture, 2 L of TB media with 25 mg/mL kanamycin was inoculated and grown at 37 °C until the OD600 reached 1.5. To induce protein expression, the culture was supplemented with 30% L-rhamnose (Sigma-Aldrich, R3875-100G) to a final concentration of 0.8%, then incubated at 18° C for 24 h with shaking. Cells were harvested by centrifugation at 4000×g for 35 min.

Cell pellets from 1 L of culture were lysed by resuspending in 30 mL of cold bacterial lysis buffer containing 20 mM HEPES, pH 7.5, 2 M NaCl, 10% glycerol, 1 mM TCEP (added fresh), and two tablets of Roche EDTA-free complete protease inhibitor cocktail. DNase I (75 U) was added, and the suspension was sonicated (10 min, 60% amplitude, 10 s on/off cycles) on ice. The lysate was clarified by centrifugation at 12,000 rpm for 30 min at 4 °C. The supernatant was incubated with 0.75 mL of pre-washed His60 nickel resin (Takara Bio) at 4 °C for 1 h to facilitate protein binding. The resin was washed sequentially with 10 mL of water and 10 mL of lysis buffer supplemented with 1 mM TCEP. After washing, the resin was loaded into a column, and the column was washed with 100 mL of wash buffer (20 mM HEPES, pH 7.5, 2 M NaCl, 10% glycerol, 1 mM TCEP, 25 mM imidazole). Proteins were eluted with 4 mL of elution buffer (20 mM HEPES, pH 7.5, 10% glycerol, 1 mM TCEP, 500 mM imidazole), collecting fractions after 10 min of incubation. To further purify the protein, the elution fractions were diluted with 50 mL of low-salt buffer (20 mM HEPES, pH 7.5, 10% glycerol, 1 mM TCEP) and loaded onto a HiTrap SP HP cation exchange column (Cytiva, 17115101) connected to FPLC system. The column was equilibrated in low-salt buffer, and proteins were separated with a gradient from 100% low-salt buffer to 80% high-salt buffer (20 mM HEPES, pH 7.5, 2 M NaCl, 10% glycerol, 1 mM TCEP) over 50 mL at a flow rate of 5 mL/min. Fractions were collected and analyzed on a 4–12% Bis-Tris gradient gel using MOPS buffer. Fractions containing the target protein were pooled and concentrated using an Amicon Ultra-0.5 Centrifugal Filter Unit (Sigma, UFC5100BK), flash-frozen, and stored at –80 °C.

### TDAC-seq integrated with CRISPR perturbation

The sgRNA sequence targeting the *HOXA7/9* CTCF site is GCCATCTGCTGGCCGCCGTT, and those for the HS2 library are listed in **Supplementary Data S1**. Oligos containing the sgRNA sequences were cloned into pLentiGuide-Puro (gift from Feng Zhang, Addgene plasmid #52953). For the pooled HS2 library, each sgRNA was pooled at equal ratios. Lentiviral particles carrying the plasmids were generated as described above and titered according to published procedures.^42^ For the HS2 screen, K562 cells (2.5 x 10^5^) were transduced at a multiplicity of infection < 0.3 and subsequently selected with puromycin. For individual sgRNA transduction experiments, virus titration was omitted. After 5 days of puromycin selection (1 µg/mL puromycin for MOLM-13 and 2 µg/mL puromycin for K562), cells were electroporated with Cas9 or ABE8e. 4 x 10^5^ cells were prepared for electroporation by washing twice in PBS, and cells were resuspended in Resuspension Buffer R at a concentration of 2 × 10^7^ cells/mL. 37.2 µM Alt-R Cas9 protein (IDT) was prepared by diluting 62 µM stock with Buffer R. For each electroporation, 1 µL of diluted Cas9 or purified ABE8e (14 µM) was mixed with 11 µL of cell suspension and electroporated at 1000 V with 50 ms pulse width for 1 pulse using the Neon Transfection System 10 µL kit (Thermo Fisher, MPK1025). Electroporation was performed twice, and cells were transferred into pre-warmed RPMI medium with 10% FBS (without antibiotics). After 6 days post-electroporation, TDAC-seq was performed and samples were sequenced as described above. For single guide experiments, 200-400 ng of gDNA was amplified in PCR1, and for the pooled screen, the amount of amplified gDNA was increased proportionally to the library size by increasing the number of PCR1 reactions.

### ATAC-seq

Cells were washed twice with cold PBS and resuspended in PBS at a density of 2 × 10^6^ /mL, ensuring a final reaction cell count of approximately 1 × 10^4^ cells per 5 µL. Next, 5 µL of cells in PBS were mixed 42.5 µL of transposition buffer. The reaction was incubated at room temperature for 10 min, followed by the addition of 2.5 µL of assembled Tn5 transposase, and then incubated at 37 °C for 30 min in a ThermoMixer set to 300 rpm. DNA was subsequently purified with the Qiagen MinElute PCR Purification Kit (Qiagen) and minimally amplified for sequencing, following established protocols.^44^ Final libraries were purified again with the MinElute PCR Purification Kit (Qiagen) and sequenced on a NextSeq 550 or Novaseq SP.

After removing adapters using NGmerge, the ATAC-seq data was aligned using bowtie2, and duplicates were removed by using picard MarkDuplicates. Bam files were converted into fragments files by using bedtools bamtobed. The genomic region of interest was divided into 1 bp bins and the number of Tn5 fragment ends overlapping with each bin was quantified as the Tn5 insertion density. Then the insertion density track was smoothed by a 250 bp window to obtain the final coverage track. The coverage tracks were exported to bigwig format and visualize in IGV.

### Quantitative reverse-transcription PCR

Cells were harvested for total RNA isolation using the RNeasy Mini Kit (Qiagen). cDNA synthesis was performed with the Applied Biosystems™ High-Capacity cDNA Reverse Transcription Kit (Applied Biosystems™ 4368814). Quantitative reverse transcription PCR (qRT-PCR) was conducted using SYBR™ Select Master Mix on the CFX Connect Real-Time PCR Detection System (Bio-Rad) with oligonucleotide primers listed in **Supplementary Data S2**. Results were calculated as the fold-change in mRNA expression of the gene of interest, normalized to GAPDH expression, using the ΔΔCt method.

### DddA-whole genome sequencing analysis

Nanopore sequencing data were aligned using Minimap2.^45,46^ Duplicate reads were removed, and base-specific mutation counts were extracted using Samtools mpileup. Mutation rates at each cytosine were calculated, saved as a bedGraph file, and converted to a BigWig file. The distribution of cytosine deamination fractions around CTCF binding sites or TSS was analyzed and plotted using computeMatrix and plotProfile from deepTools

### Detection of DddA base edits from TDAC-seq

Nanopore sequencing data were aligned using the Minimap2.^45,46^ Reads with larger than 500 bp unaligned regions compared to the full amplicon on either end of the read were removed. Single base mutations and indels were extracted using the cs tag after alignment. CpG sites were masked from mutation calling to prevent bias introduced by endogenous CpG methylation. Only C-to-T and G-to-A mutations were recorded as potential DddA edits. Reads with more C-to-T than G-to-A mutations were assigned to the top strand of the template and vice versa. TDAC-seq signals were compared with published ATAC-seq and ChIP-seq data, which are listed in **Supplementary Data S3.**

### Computing DddA sequence bias

DddA sequence bias was estimated using DddA edit rate on naked DNA with a k-mer model (k=3). k-mers with center bases being A and T were assigned with a bias of 0. 8 regions in the human genome were selected, and the DddA editing experiment was performed. Regions at chrX:48798803-48803519, chr8:127733151-127737826, chr3:141366125-141371787, and chr7:107741812-107745536 were selected for model fitting, and regions at chr1:170530445-170533865, chr21:38118238-38127665, chr5:180324459-180334556, and chr7:27158522-27163197, were selected for model testing.

### Calculating DddA footprints

DddA footprints were computed either on individual DNA molecules or aggregated reads. For each single base-pair position of interest. The sum of DddA edits were computed for a center region with a given radius *r*, as well as two flanking regions with a diameter of *r*. Similarly, the sum of DddA sequence bias was computed for the same center and flanking regions. Then the depletion of DddA edits at the center was calculated using a left-tailed center-versus-flank binomial test using bias_center / (bias_center + bias_sum) as the probability *p* of binomial distribution. A pseudo-count was added to prevent division of zero. Testing was performed for the left and right flank separately, and the least significant *p*-value was kept as the final *p*-value. Eventually, the -log10(*p*-value) was used as the footprint score. DddA footprints were compared with published MNase-seq and ChIP-seq data, which are listed in **Supplementary Data S3**.

### Analysis of pooled CRISPR screen data

CRISPR deletions were detected during alignment of nanopore sequencing reads by parsing the cs tag. Deletions larger than 5 bp were kept as CRISPR deletions. To estimate the effect of each guide, we compared wild-type reads to those with deletions at the site of interest and without any deletion outside this window were selected as foreground reads. The foreground and background reads were first down-sampled to the same read number, then deduplicated. Then the average DddA accessibility, as well as footprints were compared between the foreground and background. Here, the comparison was performed separately for the C-to-T strand and G-to-A strand and then averaged to prevent any bias introduced by imbalanced strand sampling. Additionally, DddA signal within the window of interest is masked to be 0 to prevent false positive differences resulting from the CRISPR deletion itself. For differential footprints, footprints were first calculated on single DNA molecules before aggregating to obtain the final result.

### Identification of minimal regulatory element from diverse CRISPR deletions

We considered whether diverse CRISPR deletion genotypes share a common mechanism by deleting the same minimal regulatory element. To nominate such a regulatory element, we systematically considered all 1-15 bp (deletion window) subsequences. For each candidate subsequence, we grouped reads that contain a deletion spanning the entire window and where the deletion length is at most 10 bp longer than the window. All other reads were assigned to a second group. Welch’s *t*-test was performed to compare the number of DddA edits per read in the two groups, and we used the *p*-value to reflect the quality of using the candidate deletion window to group reads, similar to traditional clustering algorithms. Windows with high purity correspond to a subsequence whose deletion is necessary and sufficient for an effect on DddA accessibility.

### Analysis of pooled ABE screen data

ABE base edits were detected during alignment of nanopore sequencing reads by parsing the cs tag. Adenine bases in positions +3 to +9 of the sgRNA were used for analysis. The ratio of editing rate at each adenine in the presence and absence of ABE treatment was used to threshold candidate adenine positions (threshold is dependent on the dataset), and the strand of A-to-G edits was examined to ensure the correct adenines were selected. Instead of comparing sgRNAs, the effect of editing at each individual adenine was estimated by comparing reads where the adenine of interest was edited with reads without any edits throughout. Then edited and unedited reads were down-sampled to the same number, deduplicated, and used to compare DddA accessibility.

### Comparison to TF binding sites

High confidence TF binding sites were obtained from Unibind.^47,48^ The individual TF binding events were superimposed onto the genomic region of interest. Overlapping TF binding site annotations were merged. The final TF binding event list was visualized using the DNA Features Viewer Python package.

### Sequencing read deduplication

To reduce the effect of PCR bias, we identified and removed sequencing reads from PCR duplicates using the pattern of DddA edits across the amplicon. Because the amplicon is large, it is unlikely for any pair of cells to receive C·G-to-T·A edits in the exact same or extremely similar positions, so such reads are attributed to PCR duplication. As such, for each read, we extracted a binary vector indicating whether each C·G in the reference has been edited to T·A and used this as a unique molecular identifier (UMI). UMIs are grouped together using UMI-tools^49^ if their hamming distance is less than 1% of the UMI length, which is approximately the error rate of Oxford Nanopore sequencing. Finally, for each group, the read with the most common UMI is kept.

### Calling deaminase-accessible DNA sequences (DADs)

For each read, we identified regions with a high density of DddA edits, termed deaminase-accessible DNA sequences (DADs). To account for the enzyme’s substrate preferences (i.e. it only edits cytosines, which naturally occur at variable densities in the genome, and its editing efficiency is strongly determined by the preceding nucleotide), we used a hidden Markov model based on the hmmlearn implementation but with custom emission probabilities (**Figure S1D**). In the model, each position is in either an open region (DAD) or a closed region. In the former case, the C·G-to-T·A editing rate is higher and depends on sequence context because such events are attributed to DddA editing. Meanwhile, in the latter case, the editing rate is lower because such events are attributed to sequencing error. The model parameters were obtained by fitting the sequencing data for each TDAC-seq locus (**Figure 1E,F, Figure S1C,E)** and selecting a representative, well-performing model.

---

**Figure S1.**
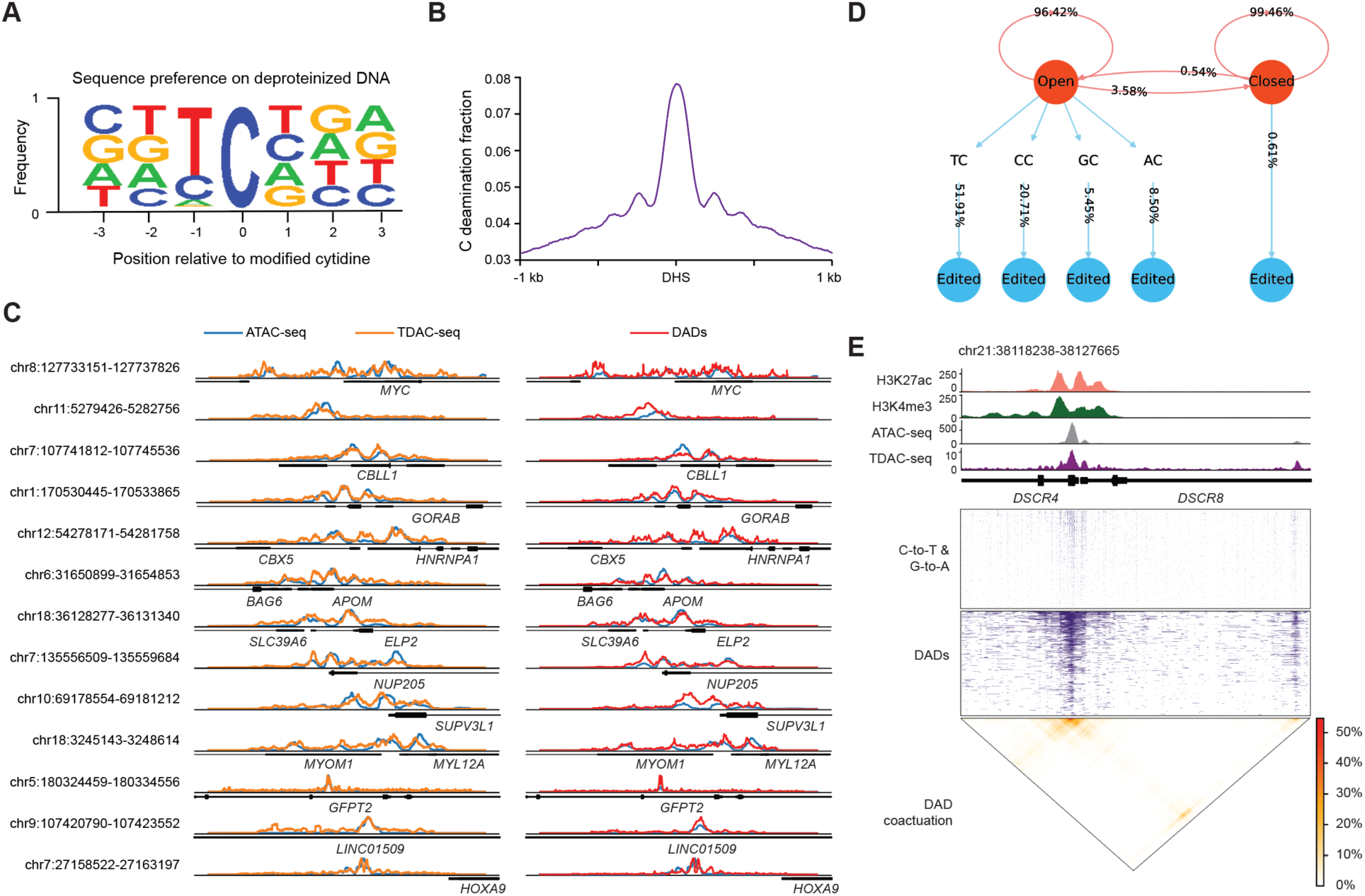
Supporting Data for Figure 1. **A.** Sequence logo showing the sequence context preference for DddA11 on naked DNA. **B.** Aggregate profile plot showing distribution of cytidine deamination fractions (*y* axis) around DNase I hypersensitive sites (DHSs) from whole-genome sequencing (WGS) of DddA11-treated K562 cells. **C.** Genome tracks showing signal from ATAC-seq (blue), TDAC-seq (orange), and DADs (red) across indicated genomic regions, indicating that aggregate TDAC-seq and DAD signals align with ATAC-seq. **D.** Schematic of hidden Markov model used to call DADs. Hidden states are orange, and observables are blue. **E.** Top: genome tracks showing chromatin profiles, TDAC-seq, and DADs identified for each single DNA molecule at the *DSCR4/8* locus. Bottom: heatmap showing DAD co-activation, where color indicates the percentage of reads where both positions are open on the same DNA molecule.

**Figure S2.**
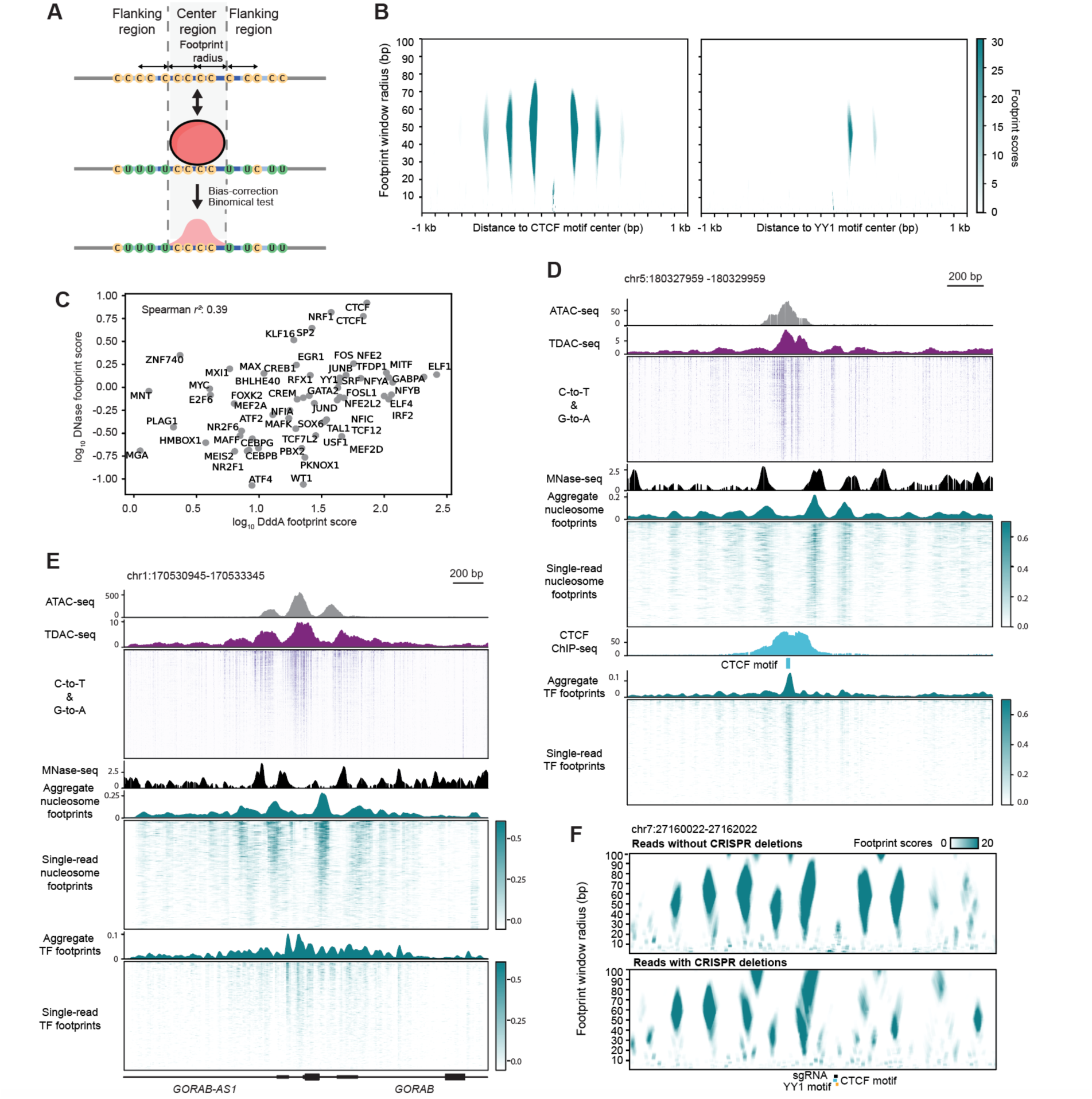
Supporting Data for Figure 2. **A.** Schematic of DddA11 footprint calling method. **B.** Heatmap showing DddA11 footprints from DddA11-WGS data centered around CTCF (left) and YY1 (right) binding sites calculated with varying footprint radii (*y* axis). **C.** Scatter plot comparing DNase-seq TF footprint scores with DddA11 footprint scores quantified for different TFs. **D.** Genome tracks showing ATAC-seq, TDAC-seq, MNase-seq, CTCF ChIP-seq and DddA11 footprint scores at indicated genomic locus in K562 cells **E.** Genome tracks showing ATAC-seq, TDAC-seq, MNase-seq, and DddA11 footprint scores at the *GORAB-AS1/GORAB* locus in K562 cells **F.** Heatmaps showing aggregate DddA11 footprints across the same reads from **Figure 2A** and Figure 2C calculated with varying footprint radii (*y* axis).

**Figure S3.**
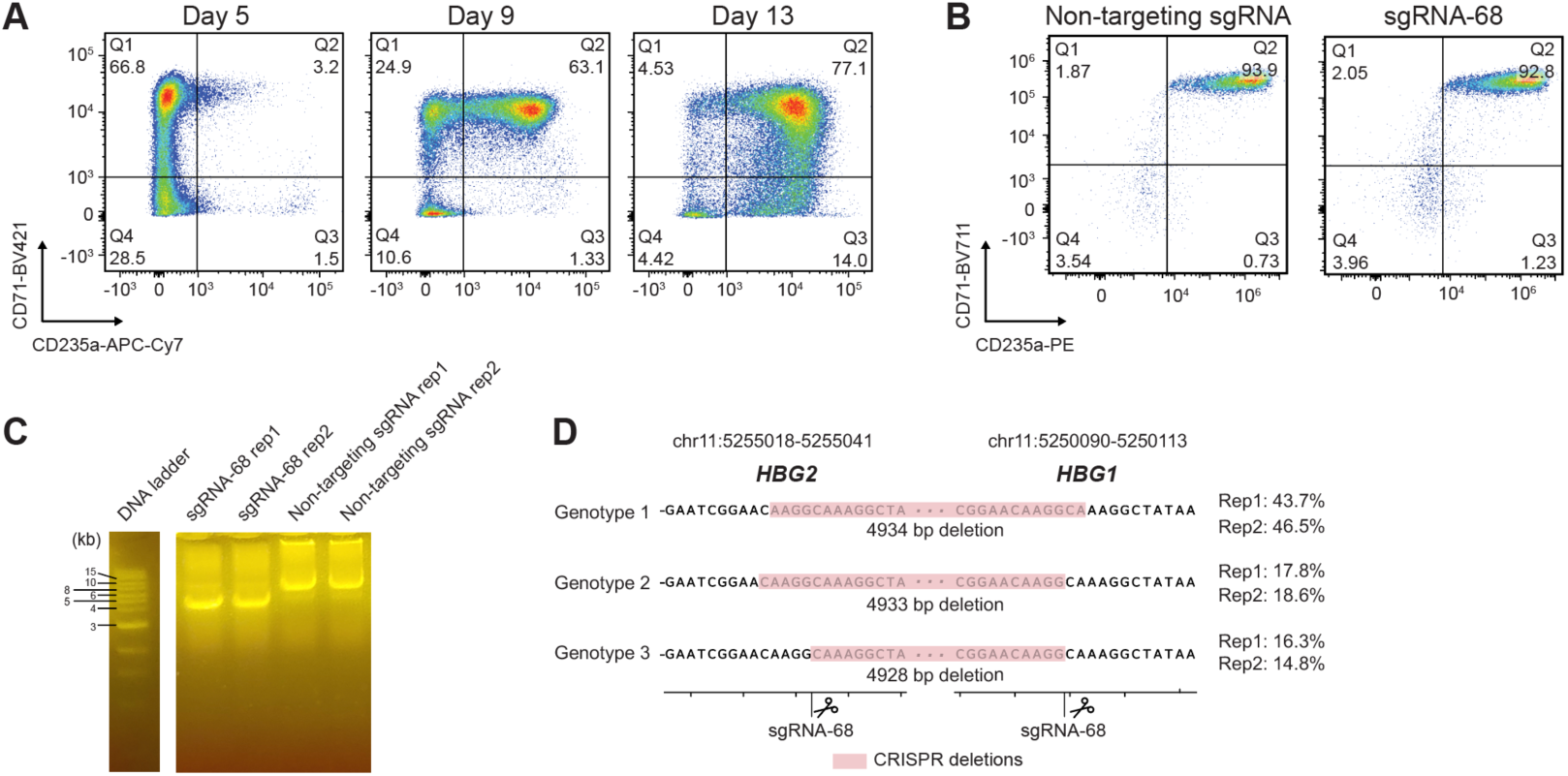
Supplementary Data for Figure 3. **A.** Flow-cytometry measurements of CD71 and CD235a levels in erythroid-differentiated HSPCs on day 5, 9 and 13. **B.** Flow-cytometry measurements of CD71 and CD235a levels in erythroid-differentiated HSPCs (day 5), transduced with indicated sgRNAs and electroporated with Cas9 protein. **C.** DNA gel of TDAC-seq library amplified at the HBG1/2 locus from erythroid-differentiated HSPCs (day 5), transduced with sgRNA-68 or non-targeting sgRNA and electroporated with Cas9 protein. **D.** Top three abundant genotypes (left) at the *HBG* locus from erythroid-differentiated HSPC cells (day 5) transduced with sgRNA-68 with their relative abundance in sequencing reads (right).

**Figure S4.**
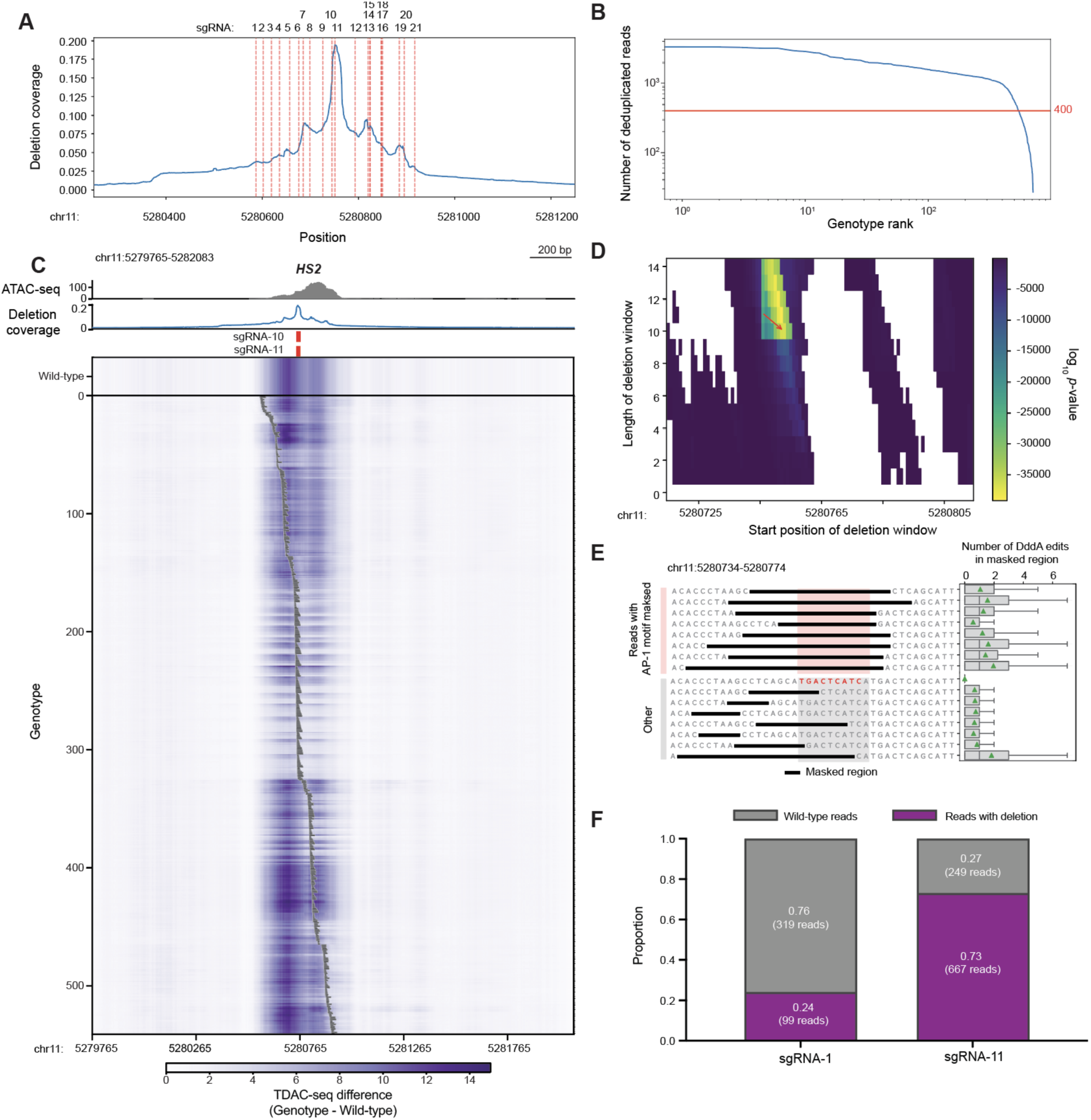
Supplementary Data for Figure 4. **A.** Line plot showing the fraction of reads (*y* axis) containing a deletion at the indicated position across the HS2 region (*x* axis), with expected cut sites for each sgRNA indicated with red dotted lines. **B.** The number of deduplicated reads (*y* axis) per genotypes (*x* axis) from HS2 CRISPR deletions. In the HS2 Cas9 TDAC-seq sgRNA tiling library, 541 genotypes have at least 400 reads (red line). **C.** Heatmaps of TDAC-seq using CRISPR-Cas9 cutting on the HS2 enhancer in K562 cells. Individual sequencing reads were assigned to one of 541 genotypes (*y* axis), with black lines indicating the CRISPR deletion position and length on the HS2 region. For each genotype, TDAC-seq signals are shown across the HS2 region (*x* axis). **D.** Heatmap showing the statistical significance (Welch’s *t*-test) of comparing DddA accessibility between reads that contain a deletion spanning the entire deletion window and all other reads. This value is calculated for all possible deletion windows indicated by their start position (horizontal axis) and length (vertical axis). Those with less than 100× coverage were excluded. Arrow indicates a 10-bp deletion window corresponding to the minimal AP-1 motif discussed in **Figure 4**. **E.** Box plot (right) showing the number of DddA edits in masked regions (black bars) from wild-type reads. The genotypes are the same as those in **Figure 4C**. The number of DddA edits in the masked regions is minimal and does not significantly impact the total DddA edits in the accessible region. Box plots show the median and interquartile range, with whiskers extending to the minimum and maximum values, excluding outliers that are beyond 1.5× the interquartile range. Mean is indicated by green triangles. **F.** The proportion of reads from gDNA with and without CRISPR deletions after individual transduction of sgRNA-1 and sgRNA-11 into K562 cells.

**Figure S5.**
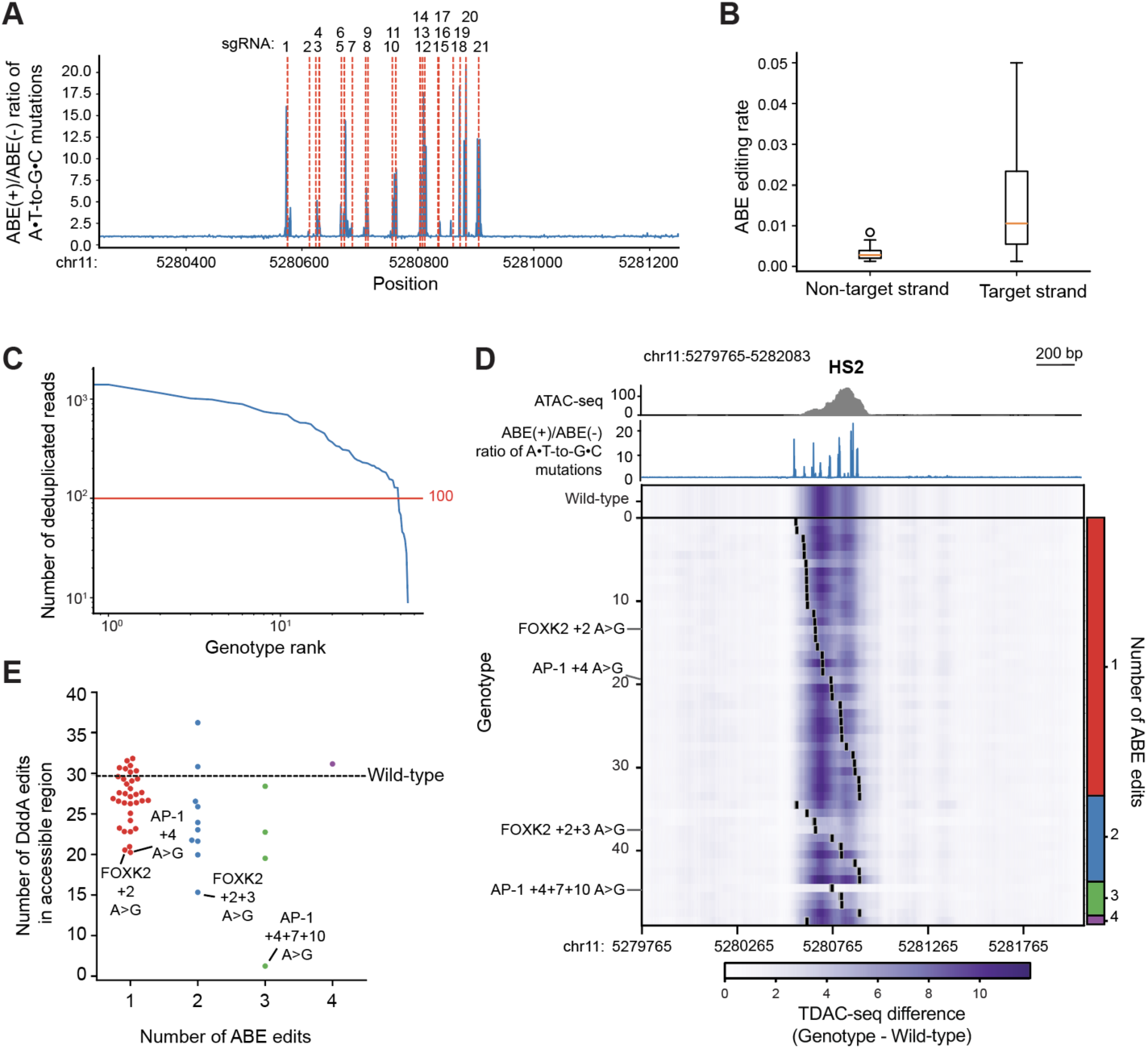
Supplementary Data for Figure 5. **A.** Line plot showing the ratio of A•T-to-G•C edit coverage in ABE(+) versus ABE(-) samples with sgRNA predicted editing sites based on the ABE8e editing window (red dotted lines) across the HS2 region (*x* axis). **B.** Box plot of A•T-to-G•C mutation rate at adenosines on the non-target strand and target strand for the HS2 pooled ABE screen. Target strand is defined as the strand containing the same sequence as the sgRNA, and non-target strand is the strand containing complementary sequence to the sgRNA. The boxes define the first quartile (Q1), median, and third quartile of data (Q3), and whiskers extend to the farthest data point within Q1 - 1.5x the inter-quartile range (IQR) and Q3 + 1.5x IQR. Data points outside of the whisker range are shown as fliers. **C.** Number of reads (*y* axis) per genotype (*x* axis) from HS2 ABE edits. In the HS2 ABE TDAC-seq sgRNA tiling library, 49 genotypes have at least 100 reads. **D.** Heatmaps of TDAC-seq integrated with pooled ABE base editing on the HS2 enhancer in K562 cells. Individual sequencing reads were assigned into one of 49 genotypes (*y* axis), with black lines indicating the A•T-to-G•C mutation sites on the HS2 region. For each genotype, TDAC-seq signals are shown across the HS2 region (*x* axis). Genotypes were grouped by the number of ABE edits (right). **E.** Swarm plots showing the number of DddA edits in the accessible region (**Figure 5A**) per genotype, stratified by the number of ABE mutations. Horizontal line represents wild-type.

